# A New Disease Gene for Hypokalemic Periodic Paralysis, *KCNA7*, Established in a Multigenerational Family

**DOI:** 10.64898/2026.09.09.750381

**Authors:** Khurram Liaqat, Gourav Saha, Shuai Guo, Kayla Treat, Fenfen Wu, Summan Thahiem, Undiagnosed Disease Network, John C Kincaid, Michael Rubart-von der Lohe, Francesco Vetrini, Stephen Cannon, Erin Conboy

## Abstract

Hypokalemic periodic paralysis (HypoPP) is an inherited skeletal muscle ion channelopathy of *CACNA1S* or *SCN4A* characterized by recurrent episodes of weakness, often triggered by rest after exercise or by reduced K+ (carbohydrate ingestion, stress). Here, we describe a multigenerational family in whom a *KCNA7* missense variant [c.834A>C (p.Arg278Ser)] co-segregated with susceptibility to recurrent attacks of weakness in association with hypokalemia as low as 1.3 mEq/L and a 40% decrement of the compound muscle action potential after exercise, suggesting an additional HypoPP gene. Arg278 is the outermost positively charged residue (R1) within the S4 transmembrane segment of the voltage-sensor domain of K_V_1.7, orthologous to the canonical patten of arginine missense mutations at R1 or R2 in S4 segments for HypoPP-mutant Ca_V_1.1 and Na_V_1.4 channels. To determine the functional consequences of Arg278Ser, we expressed wild-type and mutant K_V_1.7 channels in *Xenopus* oocytes and HEK293 cells. Arg278Ser, but not wild-type K_V_1.7, generated an anomalous inwardly rectifying current at hyperpolarized membrane potentials that was nonselective Na^+^ or K^+^, consistent with the anomalous gating pore conductance that causes susceptibility to HypoPP. Other pathogenic variants of *KCNA7* have previously been implicated only in inherited cardiac arrhythmias. Collectively, the genetic, structural and electrophysiological findings support *KCNA7* Arg278Ser as a pathogenic variant underlying HypoPP and implicate *KCNA7* as a disease gene for periodic paralysis, extending the established gating-pore mechanism of HypoPP to K_V_1.7.

## Introduction

Hypokalemic periodic paralysis (HypoPP) is an inherited ion channelopathy of skeletal muscle characterized by transient episodes of muscle weakness associated with marked reductions in serum potassium due to intracellular potassium redistribution. These episodes are often triggered by metabolic or physiological factors such as high-carbohydrate intake, rest following strenuous exercise, infection, emotional stress, or dehydration^1–3^. Symptom onset typically occurs during adolescence, although it can range from childhood to the third decade of life^4^.

The prevalence of HypoPP is estimated to be approximately 1 in 100,000 individuals^5^. Hypokalemic periodic paralysis (HypoPP) is most frequently caused by missense variants in *CACNA1S* encoding the Ca_V_1.1 channel (70% of families), but in 10% of families the HypoPP is caused by variants in *SCN4A* encoding Na_V_1.4 channel while up to 20% remain unidentified^6,7^. The overwhelming majority of established pathogenic variants associated with HypoPP cause missense mutations of Arg residues in the S4 segment^8^ (outermost R1 or R2 positions) for either Ca_V_1.1 (9 out of 10) and Na_V_1.4 (15 out of 15). This consistent pattern for the location of HypoPP missense mutations has a correspondingly conserved functional defect^9,10^. Functional expression studies have revealed an anomalous inward-rectifying current at hyperpolarized potentials for 6 HypoPP variants of Ca_V_1.1, including the non-canonical Val876Glu, and for 11 of the Na_V_1.4 variants. This so-called gating pore current contributes a small leakage current when the muscle fiber is hyperpolarized at rest (−90 mV) that causes susceptibility to paradoxical depolarization of the fiber in low K^+^ (< 3.5 mM) thereby inactivating Na^+^ channels, reducing fiber excitability, and causing ictal muscle weakness^3^.

In addition to the well-established HypoPP genes *CACNA1S* and *SCN4A*, other genes have been associated with atypical forms of periodic paralysis that share some clinical features with HypoPP. ATP1A2, encoding the α2 subunit of the Na /K -ATPase^11^, the mitochondrial genes *MT-ATP6* and *MT-ATP8*, encoding ATP synthase subunits^12^; *RYR1*, encoding ryanodine receptor type 1^13^; and MCM3AP, encoding minichromosome maintenance 3-associated protein^14^.

In this study, we identified and characterized a multigenerational family in which affected individuals had typical features HypoPP and yet did not have variants identified in a panel of 4 periodic paralysis genes (*CACNA1S, SCN4A, KCNJ2,* and *RYR1*). Further genetic studies identified a novel missense variant [c.834A>C (p.Arg278Ser)] in *KCNA7* that cosegregated with the affected family members. The *KCNA7* gene encodes K_V_1.7, a member of the Shaker subfamily of voltage-gated potassium channels highly expressed in skeletal muscle and much lower levels in heart^15^. The *KCNA7* gene has previously been reported as a candidate gene for inherited cardiac disorders^16^. The Arg278Ser missense mutation is in the R1 position of the S4 segment, homologous to established HypoPP variants of Ca_V_1.1 and Na_V_1.4. Furthermore, the functional studies using *Xenopus oocyte* expression and HEK293T cells demonstrated that the *KCNA7* variant Arg278Ser produces an abnormal inward-rectifying current at hyperpolarized potentials, consistent with a HypoPP-associated gating pore current, thereby supporting its pathogenic role, suggesting a novel mechanism of disease.

## Material and Methods

All procedures were conducted in accordance with the ethical standards of the respective institutions. This family was recruited through the Undiagnosed Rare Disease Clinic (URDC) at Indiana University. Written informed consent was obtained from all the members participating in this study for the collection, research use, and storage of the specimens in accordance with the approved protocol by the Indiana University Institutional Review Board (IRB# 2005902680).

### Genetic Testing

Prior to the family’s enrollment in the URDC, extensive genetic testing had been performed for multiple family members, including the Invitae Periodic Paralysis Gene Panel and clinical exome sequencing. The Periodic Paralysis Gene Panel analyzed the following four genes: *CACNA1S*, *KCNJ2*, *RYR1*, and *SCN4A*. Following enrollment of the family in the URDC, the research-based ES analysis was performed.

### Oocyte expression

The plasmid IDG_KCNA7_OE_1 containing the human Kv1.7 insert was a gift from Mike McMannus (Addgene #161680). The human *KCNA7* coding sequence was PCR amplified and subcloned into the pGEMHE oocyte expression vector^17^. Mutagenesis of hKv1.7 was performed using the Q5 site-directed mutagenesis kit (New England Biolabs) to create Arg278Ser and verified with Plasmidasaurus Whole Plasmid Sequencing (Oxford Nanopore, R10.4.1). Complementary RNA was synthesized by *in vitro* transcription (2 ng/nl) with the mMESSAGE MACHINE kit (Invitrogen). Oocytes (Xenopus 1) were injected with WT, Arg278Ser, or a 1:1 mixture of RNAs (50 nl) and 1 to 3 days later ionic currents were recorded by two-electrode voltage-clamp. The microelectrodes were filled with 3 M KCl, and the external bath solution was either Na-based (in mM 100 NaCl, 1 MgCl_2_, 1.8 CaCl_2_, 10 HEPES, pH 7.1), K-based (in mM 100 KCl, 1 MgCl_2_, 1.8 CaCl_2_, 10 HEPES, pH 7.1), or N-methyl-D-glucamine (in mM 100 NMDG, 4 Ca acetate, 10 HEPES, pH 7.1 with methanethiosulfonate). Voltage steps were imposed from a holding potential of –100 mV mV and spanned the range from –150 mV to +20 mV. To optimize the detection of inward rectifying gating pore leakage currents, the background leak was determined from a linear regression on the responses from -60 mV to -45 mV and then subtracted from the measured ionic current over the entire range of voltage steps.

### Plasmid Construction and HEK293 cell expression

The expression plasmids encoding wild-type and mutant *KCNA7* were generated using the mammalian expression vector pcDNA3.4 (Addgene plasmid #253816) as the backbone. DNA inserts were cloned into the XbaI and EcoRV restriction sites of the vector. The inserts consisted of the KCNA7 wild-type sequence (KCNA7_WT; NCBI RefSeq accession no. NM_031886.3) or the corresponding KCNA7 mutant sequence [c.834A>C (p.Arg278Ser)] followed by an internal ribosome entry site (IRES) derived from encephalomyocarditis virus (EMCV) and an enhanced green fluorescent protein (EGFP) reporter sequence. The complete DNA constructs were commercially synthesized and cloned by GenScript, Inc. Following synthesis and cloning, the plasmid constructs were verified by DNA sequencing to confirm the identity and integrity of the inserted sequences. Purified plasmids were subsequently used for transfection experiments. The final constructs are herein referred to as pcDNA3.4-KCNA7_WT-IRES-EGFP and pcDNA3.4-KCNA7_Arg278Ser-IRES-EGFP. Human embryonic kidney (HEK) 293 cells (American Type Culture Collection, Manassas, VA, United States) were cultured in Iscove’s modified Dulbecco’s medium (ThermoFisher Scientific, Waltham, MA, United States) supplemented with 10% FBS and 1% penicillin-streptomycin in a 95% O_2_-5% CO_2_ incubator at 37° C. HEK 293 cells were transfected with 1μg of a plasmid encoding human wildtype (WT) *KCNA7* or Arg278Ser mutant *KCNA7* and internal ribosome entry site-enhanced green fluorescent protein (IRESeGFP), as described above, using Effectene Transfection Reagent (Qiagen, Germantown, MD, United States). Cells expressing KCNA7 were identified by virtue of their green fluorescence and used for patch-clamp experiments.

### Electrophysiology

Whole-cell current was recorded from HEK293 cells at room temperature using the patch-clamp technique in the ruptured-patch configuration. The extracellular solution contained (in mmol/L) 140 NaCl, 1.5 CaCl_2_, 1 MgCl_2_, 4 KCl, 10 HEPES, and 10 glucose (pH 7.4). The internal solution contained (in mmol/L) 144 KCl, 1.15 MgCl_2_, 1 EGTA, 10 HEPES, and 2 MgATP (pH 7.25). Pipette resistances ranged from 3 to 5 MΩ when filled with the pipette solution. After a gigaohm seal had been achieved, the test pulse current was nulled by adjusting the pipette capacitance compensation. After break in, the whole cell charging transient was minimized by adjusting whole-cell capacitance and series resistance. Sampling rate was 10 kHz and filter frequency was 2 kHz.

### 3D modeling of missense KCNA7 variant

The sequence of the wild-type protein (Q96RP8) was acquired from the UniProt databank. The sequence was mutated via text editor. Models were established using SWISS-MODEL^18^ through target template alignment tool^19^. We used the AlphaFold 3.0 computation model (AF-Q96RP8-F1) as the reference for the modeling. The quality of the modeled structures was checked in the SWISS-MODEL server using the MolProbity tool^20^ and the QMEAN scoring^19^. We have also assessed the model quality using PROCHECK^21^. Ramachandran plots were generated via PROCHECK. Polar contacts were measured using PyMol with the cutoff=3.6 and big cutoff=4.0. Stability change was predicted through DynaMut2^22^.

## Results

### Clinical features of a HypoPP family

A 54-year-old male (proband) presents to the Undiagnosed Rare Disease Clinic (URDC) for an initial enrollment visit due to longstanding history consistent with HOKPP, with symptom onset at age 8 years and a strong autosomal dominant family history spanning multiple generations. Attacks are characterized by episodic flaccid weakness involving both proximal and distal limb musculature, as well as truncal muscles, often severe enough to impair bed mobility. There is no persistent interictal weakness, and prior neurologic examinations have demonstrated normal baseline strength, suggesting preserved muscle integrity between episodes without clear evidence of fixed myopathy currently. A similar pattern of episodic weakness occurred for both males and females in four generations of the family (Fig. 1A). While both males and females are affected in this family, there is evidence of variable expressivity potentially influenced by sex-specific or hormonal factors. The proband’s daughter reported exacerbation of attacks during cancer treatment and improvement following pregnancy.

**Figure 1.**
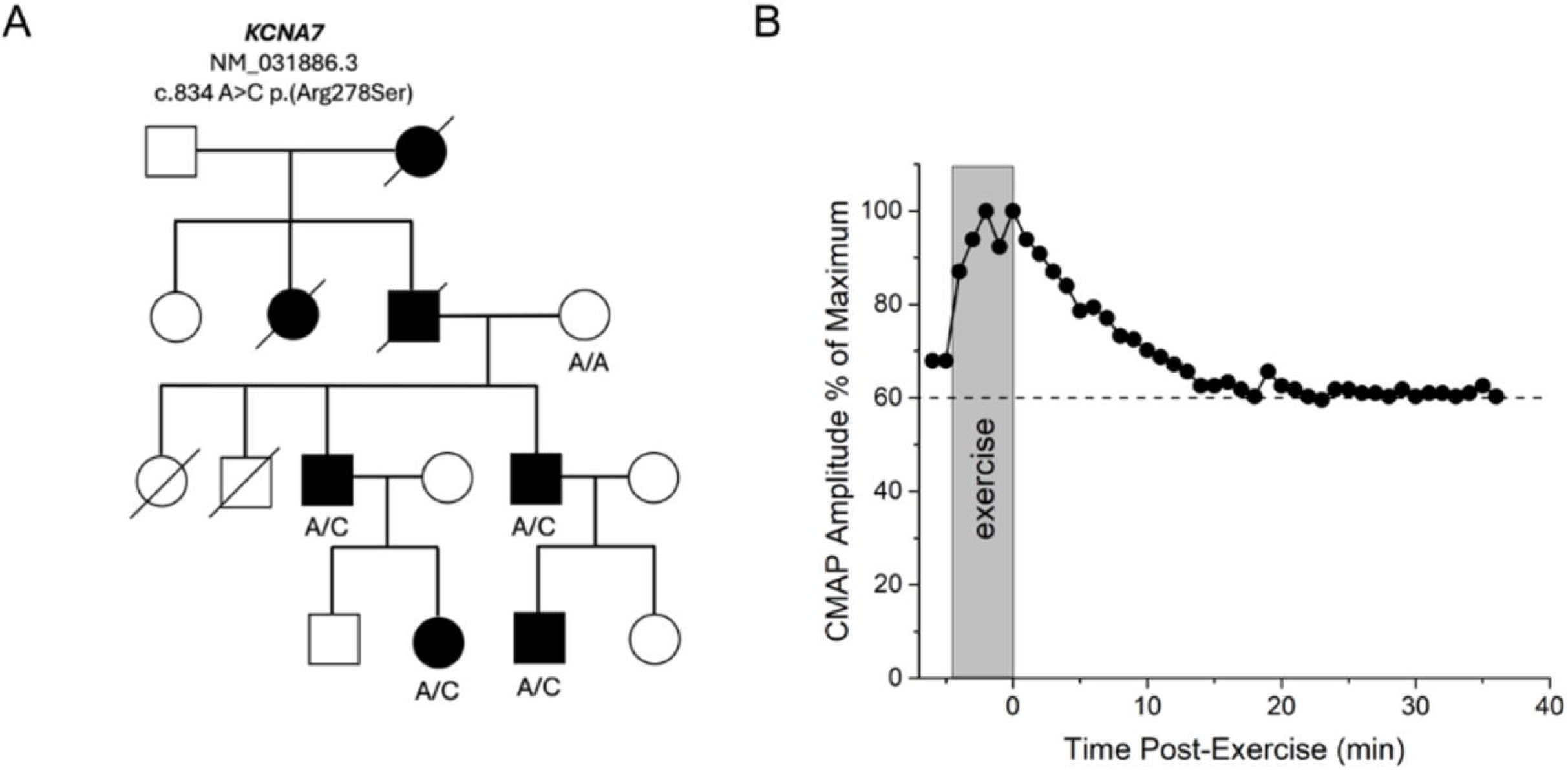
Segregation of the *KCNA7* variant and CMAP long exercise test for the proband. **(A)** Pedigree of the family showing segregation of the *KCNA7* (NM_031886.3) variant [c.834A>C; (p.Arg278Ser)]. Arrow indicates the proband. **(B)** The CMAP amplitude, relative to the maximum value during the entire test is shown before, during, and after exercise.

Attacks occur with consistent and reproducible triggers, including carbohydrate-rich intake (particularly starchy foods and sugars), emotional or physiologic stress, and overexertion. Environmental factors, notably seasonal changes and abrupt weather fluctuations, also increase attack frequency across affected family members. In contrast, exercise itself does not reliably precipitate episodes in the proband, and unlike some relatives, he does not report mitigation with light activity. Notably, his daughter describes the ability to abort or attenuate attacks with ambulation during the prodrome. Attacks demonstrate a characteristic circadian pattern, most commonly arising in the middle of the night or upon awakening. The proband endorses a recognizable prodromal “aura,” allowing early intervention with potassium supplementation and rest.

Clinical severity is variable but can be profound. During attacks, the proband is described as “floppy” and completely dependent on assistance for activities of daily living. Episode duration ranges from 30 minutes to several hours, although more severe events may persist for 24 hours to as long as 1–2 weeks. Family members demonstrate a spectrum of severity; for example, his daughter requires approximately one hospitalization annually, while his nephew reports less frequent but still clinically significant events. The proband reports an increase in attack frequency over time; however, he currently denies fixed weakness between episodes. There is no clear evidence yet of late-onset permanent myopathy, which in HypoPP may manifest as progressive weakness and, in severe cases, necessitate assistive devices such as walkers or wheelchairs.

In addition to weakness, the proband experiences significant myalgias during attacks, requiring adjunctive measures such as mechanical massage for symptomatic relief. He also reports cardiac manifestations during episodes, including irregular heart rhythms documented on electrocardiogram, a finding corroborated by other affected relatives.

The proband reports a consistent response to oral potassium supplementation, with onset of effect approximately 30 minutes after administration, though with diminishing returns beyond three doses per day. Across the family, potassium supplementation remains the primary acute management strategy, though dosing requirements vary considerably (e.g., one affected relative requires up to 8–9 doses per day). No affected family members report clinical benefits from carbonic anhydrase inhibitors.

### Clinical Laboratory Findings

Objective clinical data include documented hypokalemia during attacks and intermittent arrhythmias observed on EKG in the emergency setting. One family member had potassium levels as low as 1.3 mmol/L during severe episodes. A CMAP long exercise test performed on the proband at age 57 revealed a 40% decrement after exercise (Fig. 1B). All the available family members, including one unaffected and four affected family members, were enrolled in this study. (Fig. 1A).

### Variant identification and i*n silico* analysis

The proband was enrolled in the Undiagnosed Rare Disease Clinic (URDC) at Indiana University School of Medicine (IUSM) because a prior molecular diagnosis had not been established. The initial genetic testing, including the Invitae Periodic Paralysis panel (*CACNA1S*, *KCNJ2*, *RYR1*, *SCN4A*) and clinical exome sequencing on four affected family members was negative. Following enrollment in the URDC, research-based exome sequencing identified a heterozygous missense variant c.834A>C, (p.Arg278Ser) in *KCNA7*, in all affected individuals. *KCNA7* is highly expressed in skeletal muscle and heart^15^ and encodes a voltage-activated K^+^ channel (K_V_1.7) that slowly inactivates over hundreds of milliseconds^23^. The Arg278Ser variant, located in the S4 voltage-sensing transmembrane domain (Fig. 2A), is absent from gnomAD (v3.1 and v4.1) and is observed at a very low frequency (0.000004) in the *All of Us* dataset. It has a high CADD C-score of 21, REVEL score 0.846 and is predicted to be deleterious by various *in silico* tools including SIFT, Polyphen-2, MutationTaster. The Arginine at amino acid 278 position is highly conserved across various paralogs of *KCNA7* (Fig. 2B) and in various species (Fig. 2C).

**Figure 2.**
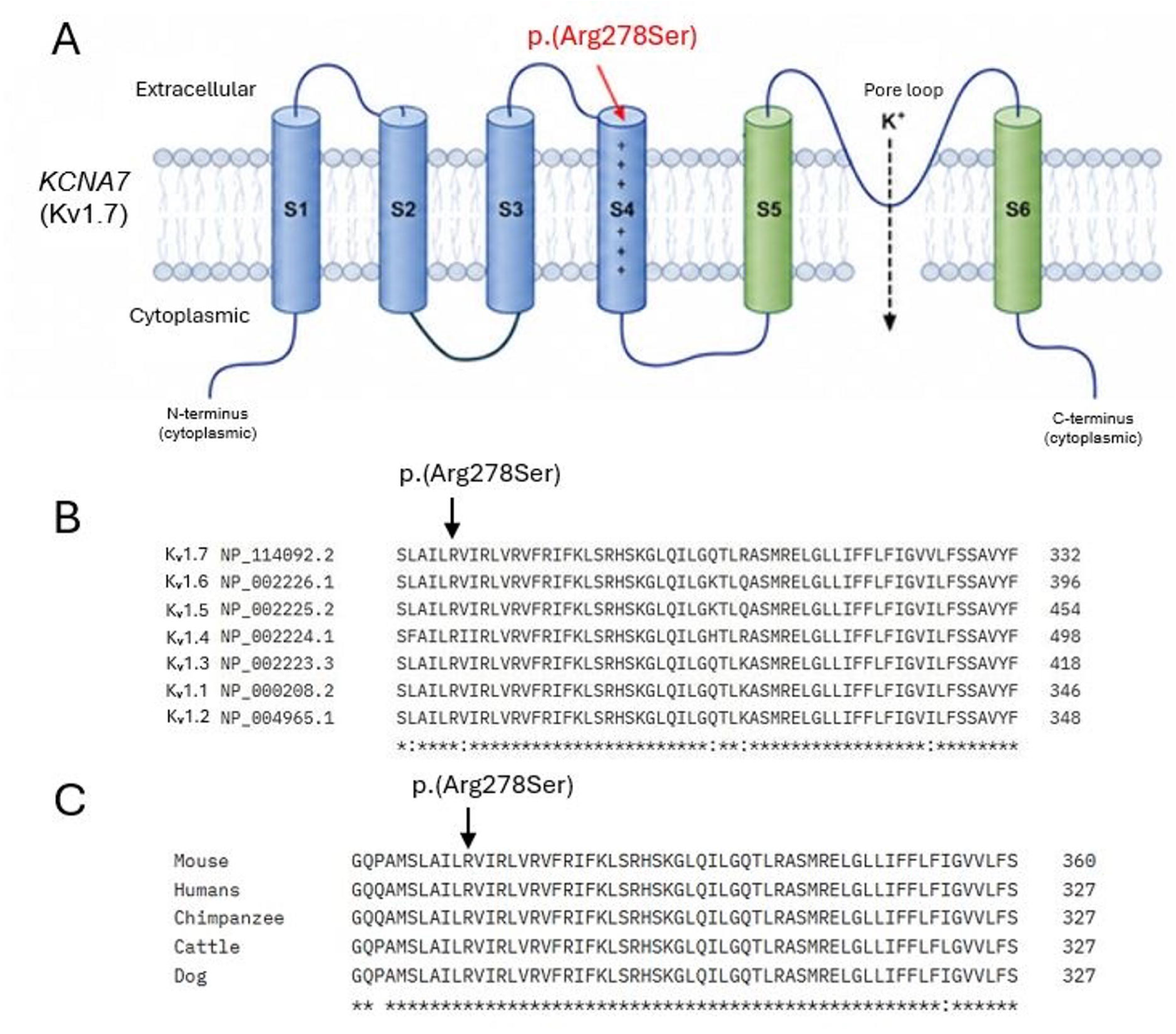
Location and conservation of Arg278 in K_V_1.7. **(A)** Schematic representation of K_V_1.7 channel topology, showing six transmembrane segments (S1–S6), cytoplasmic N- and C-termini, and the pore-loop region between S5 and S6. The Arg278Ser variant is located within the S4 voltage-sensing segment. **(B)** Multiple sequence alignment of KCNA family paralogs demonstrating conservation of the arginine residue corresponding to *KCNA7* Arg278 within the S4 voltage-sensing region. **(C)** Orthologous sequence alignment showing conservation of the Arg278 residue across representative vertebrate *KCNA7* orthologs. The red arrow indicates the position of the Arg278Ser variant.

### Functional Expression Studies

The Arg278Ser variant is homologous to the canonical pattern of HypoPP missense mutations in Ca_V_1.1 and in Na_V_1.4^3^, wherein the missense mutation of an outer arginine in S4 causes an anomalous inward rectifying leakage current, called the gating pore current. Moreover, the equivalent substitution in the Shaker K^+^ channel (Arg362Ser), an ortholog of human K_V_1.3, is known to conduct a robust gating pore current^24^. We tested whether the Arg278Ser variant identified in this HypoPP family conducts the anomalous gating pore current by recording ionic currents from oocytes injected with mRNA encoding WT or Arg278Ser hK_V_1.7. Representative currents elicited by voltage steps from -150 mV to 20 mV are shown in Figure 3. Rapidly activating outward currents were observed at test potentials >-20 mV from oocytes injected with WT or Arg278Ser mRNA. These responses are consistent with the K^+^ current conducted by the conventional pore of K_V_1.7^23^. For the Arg278Ser variant, but not WT, inward currents were observed for voltage steps of -70 mV and more negative (Fig. 3A and 3C, arrows). This inward current was observed for oocytes expressing Arg278Ser bathed in a Na^+^ solution or in a K^+^ solution. The inward currents were not detected, however, in an N-methyl-D-glucamine bath solution (Figure S2). The absence of detectable current implies the Arg278Ser leak is not permeable to a bulky organic cation, NMDG, nor is the current carried by protons as has been observed for some HypoPP Arg to His mutations in NaV1.4 channels^25^. This inwardly rectifying current that is non-selective for monovalent cations (K^+^ or Na^+^) but impermeant to NMGD is typical of gating pore currents observed in voltage-gated ion channels with missense mutations at arginines in the S4 segment of the voltage sensor domain.

**Figure 3.**
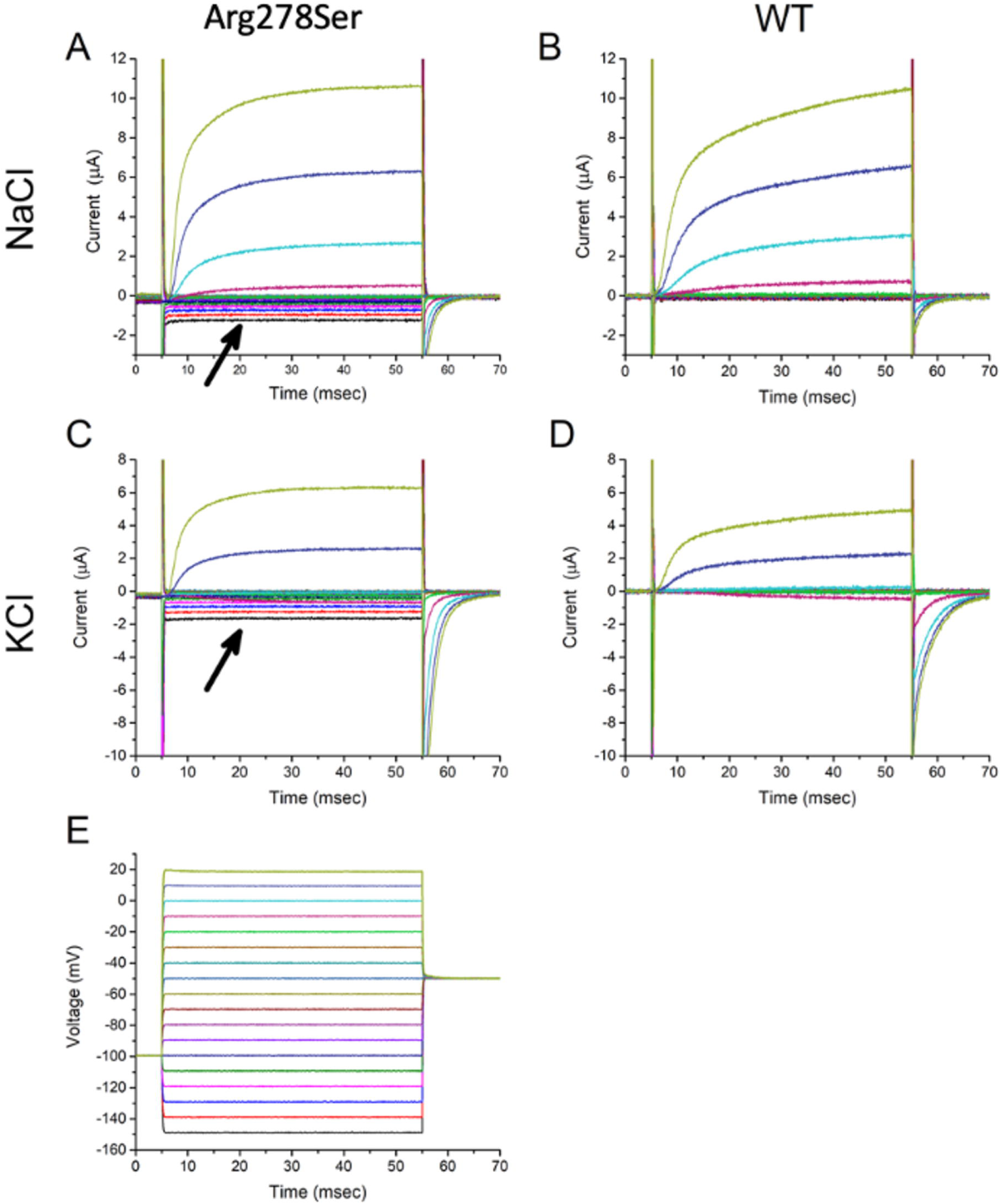
Ionic currents in oocytes injected with WT or Arg278Ser K_V_1.7 mRNA. Representative currents elicited by a 50 msec voltage step from a holding value of -100 mV are shown for an oocyte injected with Arg278Ser **(A and C)** or WT **(B and D)** mRNA. The anomalous inward current (arrows) was observed for an Arg278Ser oocyte bathed in NaCl **(A)** or KCl **(C)** but not WT oocytes. **(E)** Voltage step protocol, as recorded from the two-electrode voltage clamp of the oocyte.

The amplitude of the steady-state ionic current measured over a series of test potentials is shown in Figure 4. To combine data across multiple oocytes with variability in the peak current amplitude, the data from each oocyte were amplitude-normalized to the outward K^+^ current measured at a test potential of 0 mV in the NaCl bath. The outward rectifying current for test potentials > -20 mV was comparable for WT and Arg278Ser channels, and the reversal potential shifted to approximately 0 mV in the KCl bath (Fig. 4B), consistent with the conventional K-selective current of K_V_1.7 channels. For the Arg278Ser variant, an additional component, the gating pore current, was observed as inward rectification at voltages < -80 mV.

**Figure 4.**
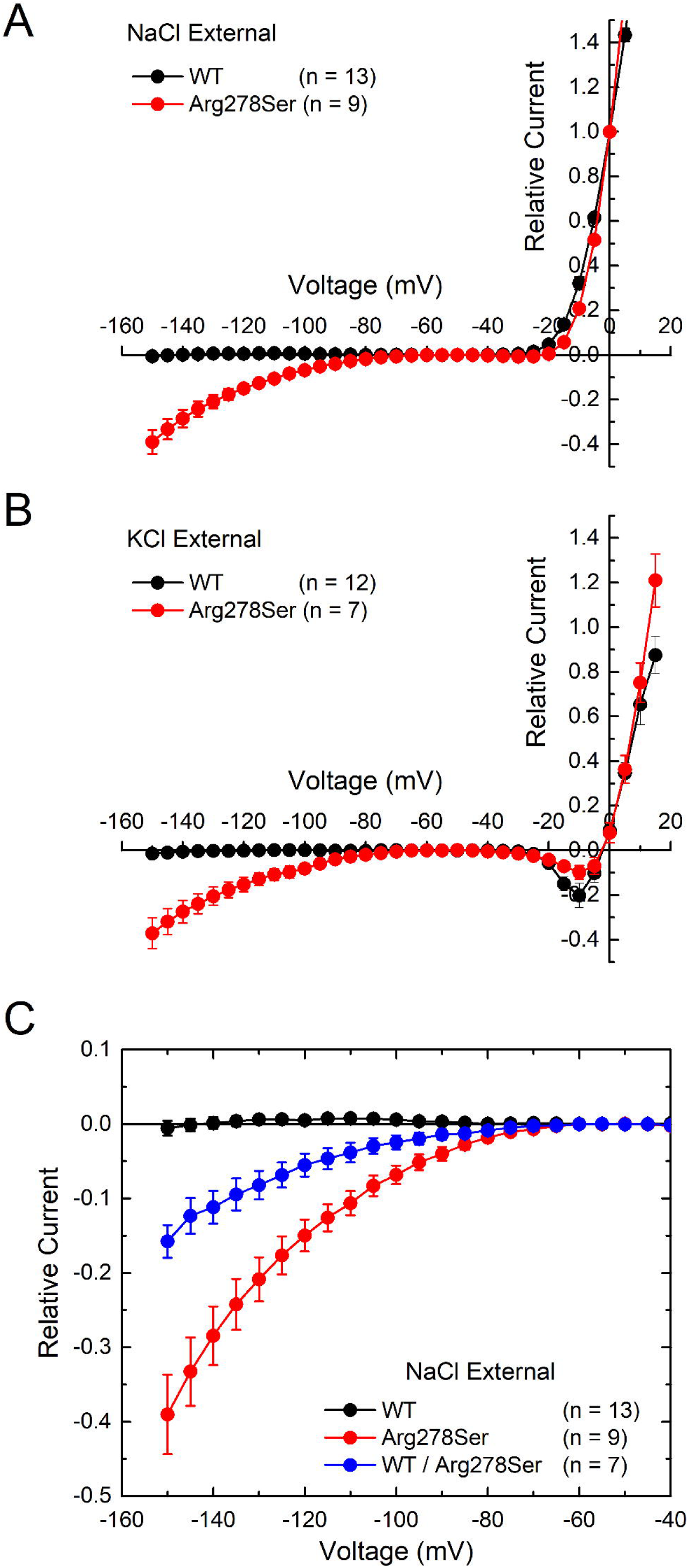
Steady-state current – voltage relationship for oocytes expressing WT or Arg278Ser hK_V_1.7 channels. Currents measured at the end of a 50 msec voltage step were amplitude-normalized to the outward current measured at 0 mV in the NaCl bath. This NaCl-based normalization, as an index of channel expression level, was used to amplitude-normalize currents measured from the same egg in a KCl bath. Inward rectifying gating pore currents were observed at voltages < -70 mV for Arg278Ser oocytes in NaCl **(A)** and in KCl **(B)** but not for WT oocytes. **(C)** Oocytes injected with a 1:1 mixture of WT and Arg278Ser mRNA had gating pore currents that were half the amplitude of 100% Arg278Ser. Data points show mean ± SEM.

Affected individuals were heterozygous for the Arg278Ser allele and so we examined ionic currents when WT and Arg278Ser were co-expressed in the same oocyte. For oocytes injected with a 1:1 ratio of WT and Arg278Ser mRNA, the amplitude of the inward rectifying gating pore current was approximately half of that observed when the same total amount of mRNA, as 100% Arg278Ser, was injected (Fig. 4C). This proportionate scaling implies the expression level of WT and Arg278Ser K_V_1.7 subunits at the plasma membrane were comparable.

We next examined whether Arg278Ser K_V_1.7 channels conduct an anomalous gating pore current when expressed in a mammalian cell line (HEK293 cells). Exemplar whole-cell currents evoked by voltage steps ranging from -160 mV to +50 mV from a holding potential of -100 mV are depicted in Figure 5A. Rapidly activating outward currents were seen at test potentials >-20 mV in cells expressing WT or Arg278Ser mutant channels. Inward currents were observed for the mutant variant, but not WT, at test potentials <-20 mV. Steady-state current-voltage relations derived from the traces shown in Figure 5A display an outwardly rectifying current component at test potentials >-20 mV for both the WT and the Arg278Ser variant, whereas an additional inwardly rectifying current component was observed for the Arg278Ser variant, but not WT, at voltages <-20 mV. Overall, average steady-state current density-voltage plots confirmed the absence of this inwardly rectifying component for the WT channel, whereas the average magnitude of the outwardly rectifying current component was significantly smaller in cells expressing mutant vs. WT channels (Fig. 5C). Control studies in untransfected HEK293 cells revealed the presence of small, rapidly activating outward currents at test voltages >-20 mV, while no significant inward currents were observed at negative test potentials (Figure S2). These responses are consistent with the voltage-gated K^+^ currents conducted by endogenous Kv channels in native HEK293 cells^26^. Overall, the dual outward and inward rectifying behavior of the current conducted by Arg278Ser K_V_1.7 channels expressed in HEK293 cells closely resembles that seen in oocytes injected with Arg278Ser K_V_1.7 RNA, suggesting that the inward rectification reflects a gating pore current of the mutant channel isoform.

**Figure 5.**
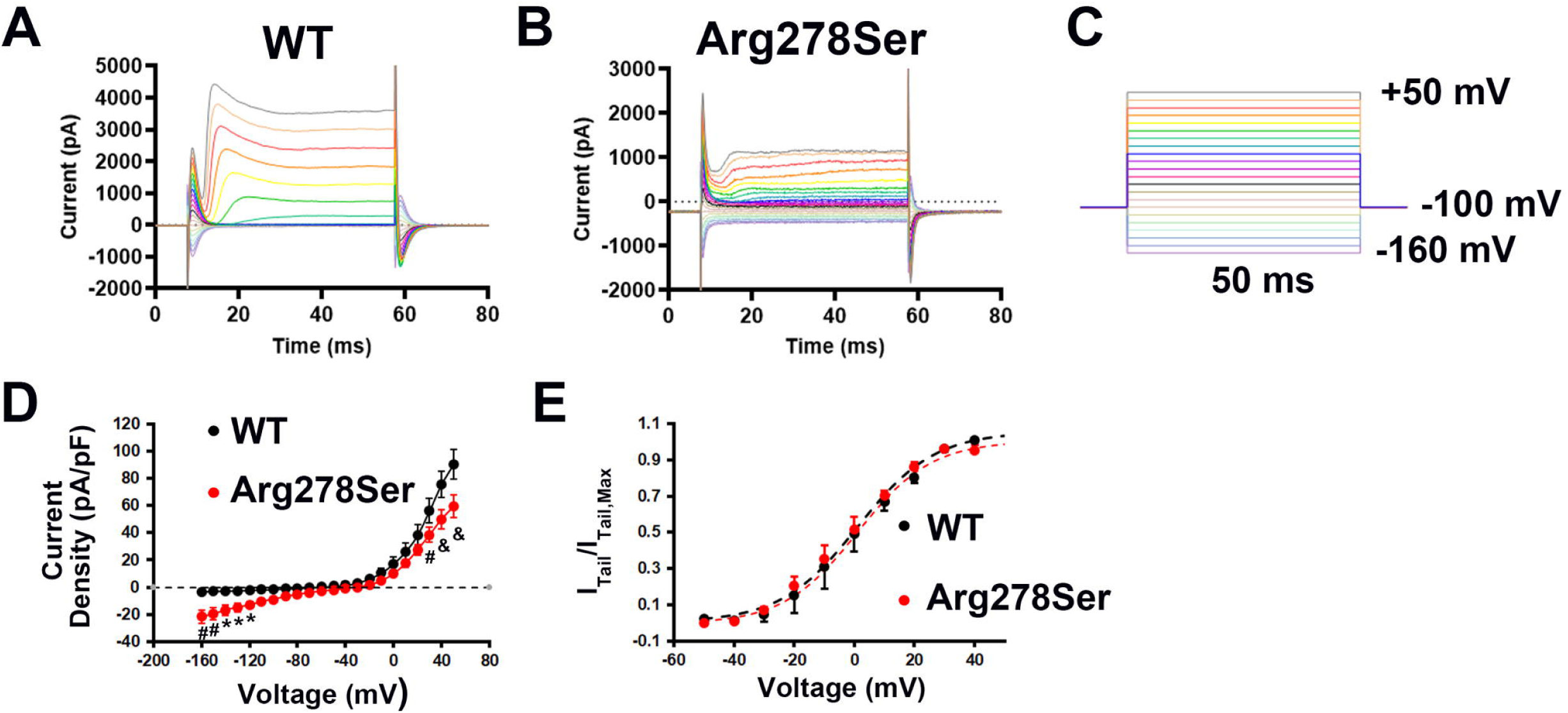
Ionic currents in HEK293 cells expressing WT or Arg278Ser K_V_1.7. Representative whole-cell current recordings for HEK293 cells expressing WT **(A)** or Arg278Ser mutant channels (**B**). Currents were elicited by 50-ms voltage commands from a holding potential of - 100 mV (**C**). Interval between voltage commands was 2 s. Traces are representative of 12 WT- and 15 Arg278Ser-expressing cells. (**D**) Steady-state current density – voltage relations for the HEK293 cell recordings. The HEK293 cell expressing Arg278Ser channels exhibit significant inward currents at voltages <-20 mV that are not seen in the cell expressing WT channels. Currents were averaged across the terminal 10 ms of each 50-ms pulse and plotted as a function of command voltage. Currents were not corrected for unspecific leak. Shown are mean ± SEM values from 12 WT and 15 Arg278Ser channel - expressing cells. *, ^#^, ^&^ *P* < 0.05, < 0.01, < 0.0001 by 2-way ANOVA and Tukey’s multiple comparisons test. (**E**) Voltage-dependence of K_V_1.7 channel activation. Average peak outward tail current density normalized to the maximal peak outward tail current density (I_Tail_/I_Tail_,_Max_) as a function of pre-pulse potential. Dashed lines represent best-fits to a Boltzmann function. Neither mid-activation voltage (WT: 5.35 [-4.8; 8.7] mV, Arg278Ser: -0.75 [-9.1; 5.5] mV; median and interquartile range; *P* = 0.29) nor slopes (WT: 9.93 [8.3; 10.5] mV, Arg278Ser: 11.1 [8.5; 13.3] mV; *P* = 0.37) were significantly different between genotypes; 6 cells per genotype; *P* values were calculated using Mann-Whitney test. Symbols and vertical lines show mean and SEM values, respectively.

Mutations in the S4 segment can shift the voltage-dependence of channel activation. Accordingly, we next assessed steady-state activation from plots of tail current amplitudes as a function of conditioning prepulse potential (Fig. 5D). These measurements revealed that the Arg278Ser mutation does not significantly alter voltage-dependence of steady-state activation of K_V_1.7 channels expressed in HEK293 cells (Fig. 5E).

## Discussion

In this study, we identified a novel *KCNA7* variant, c.834A>C (p.Arg278Ser), that segregates with autosomal dominant HypoPP in a multigenerational family. The family’s phenotype is highly consistent with primary HypoPP^1^, including childhood onset, recurrent episodes of flaccid weakness, attacks precipitated by carbohydrate-rich meals and rest or physiologic stress, documented ictal hypokalemia, preserved interictal strength, (absence of myotonia, and CMAP decrement on the long exercise test. Despite the high confidence level for a clinical diagnosis of HypoPP, a molecular diagnosis was not identified by targeted testing of canonical periodic paralysis genes. A more extensive exome sequencing screen revealed an ultrarare *KCNA7* variant, c.834A>C (p.Arg278Ser), that is not present in the gnomAD4.1 population databases, localizes to a HypoPP consensus site at R1 in the S4 segment of the voltage-sensor domain, and is predicted to be pathogenic by disrupting a conserved arginine residue (Supplementary Table 1). The expression studies demonstrated an anomalous gating-pore current in both Xenopus oocytes and HEK cells, thereby providing strong functional evidence for the pathogenicity of Arg278Ser. The importance of this discovery is bidirectional: (i) the clinical, genetic, and expression studies provide convincing evidence for a new HypoPP gene, (2) the observation of a robust gating pore current adds further to the consensus view that this anomalous inward leakage current at the resting membrane potential is the key functional defect to cause susceptibility to attacks of paradoxical fiber depolarization and weakness in HypoPP. The pathogenic current may arise from missense mutations of arginines in homologous sites of S4 segments in Ca_V_1.1, Na_V_1.4, or K_V_1.7 channels that will all result in a final common clinical phenotype of HypoPP.

HypoPP-associated variants in *CACNA1S* and *SCN4A* predominantly affect arginine residues within S4 voltage-sensing segments, supporting the concept that loss of positive charge in the voltage sensor contributes to disease pathogenesis^8^. These positively charged residues are essential for voltage sensing, and their substitution can create an anomalous gating-pore current through the voltage sensor rather than through the canonical ion-conducting pore. Such currents destabilize the skeletal muscle resting membrane potential, particularly under conditions of low extracellular potassium, leading to sarcolemmal paradoxical depolarization, sodium channel inactivation, reduced muscle fiber excitability, and episodic flaccid weakness. Consistent with this mechanism, most pathogenic *CACNA1S* variants and all of the pathogenic *SCN4A* variants identified to date involve arginine residues located in the extracellular end of S4 segments, resulting in substitution of highly conserved, positively charged arginine residues with less positively charged or neutral amino acids^27,28^. The *KCNA7* missense variant Arg278Ser identified in our family similarly replaces the outermost positively charged arginine within the S4 voltage-sensing region of K_V_1.7 with a neutral serine residue. Independent voltage-clamp studies of Arg273Ser mutant channels expressed in non-mammalian (*Xenopus* oocyte) and mammalian (HEK293) cells confirmed the presence of an anomalous gating pore conductance activated at hyperpolarized potentials (< -70 mV). Indeed, the study that first established the gating pore leakage “omega current” was for missense mutations of the outermost arginine (Arg362) in the Shaker K_V_1.1 channel^24^, a homolog of K_V_1.7. Substitution of Shaker Arg362 by Ala, Cys, His, Val, or Ser all caused the gating pore leak, with Ser resulting in the largest current. Arg278 lies within the S4 voltage-sensing helix, a region enriched for positively charged residues that move in response to changes in membrane potential. In the wild-type model, Arg278 participates in local stabilizing interactions with neighboring residues, including Leu274, Ala275, Arg281, and Leu282. Substitution with serine is predicted to reduce local electrostatic interactions, alter hydrogen-bonding geometry, and modestly destabilize the local structure. While computational modeling alone cannot establish pathogenicity, the predicted disruption is directionally consistent with the experimental finding of an abnormal leak current and with the known vulnerability of S4 arginine residues in HypoPP.

Studies in oocytes injected with WT and mutant RNA at a 1:1 ratio, mimicking heterozygosity for the mutant allele in human carriers of the mutation, still gave rise to sizable inward leak currents at negative potentials, further strengthening its pathogenicity. Results from our HEK293 cell studies do not support a role of altered voltage-dependence of activation in contributing to Arg278Ser K_V_1.7 channelopathy. However, whether changes in voltage-dependence of inactivation or recovery from inactivation occur warrants further examination.

This study has several limitations. First, the evidence is derived from a single family, and independent confirmation in unrelated individuals or families with *KCNA7* variants will be essential to establish the direct causality of the variant and the gene-disease relationship. The functional in vitro studies establish the existence of an anomalous gating pore current but may not fully recapitulate the channel behavior in human skeletal muscle. Additional studies are needed to evaluate *KCNA7* expression and localization in skeletal muscle in order to assess the magnitude of the specific conductance for the gating pore leak caused by Arg278Ser, relative to the normal ionic conductances in resting fibers. Finally, longitudinal follow-up will be important to determine whether affected individuals develop late-onset fixed myopathy, as can occur in other forms of HypoPP.

In conclusion, we have identified the *KCNA7* Arg278Ser as a pathogenic variant underlying autosomal dominant HypoPP in a multigenerational family. The variant affects a conserved S4 arginine residue, segregates with disease, is extremely rare in population databases, is predicted to perturb the local voltage-sensor structure and produces an anomalous hyperpolarization-activated inward current consistent with a pathogenic gating-pore mechanism in different experimental systems. These findings expand the genetic landscape of HypoPP and implicate K_V_1.7 as a new channel involved in skeletal muscle excitability and its perturbation in a new form of HypoPP. Additional families and functional studies are warranted to ascertain the prevalence of *KCNA7* as a recurrent HypoPP disease gene and to define its broader clinical spectrum.

## Data availability

All data supporting the findings of this study are available to those eligible upon request to the corresponding author.

## Acknowledgements

We thank the referring physicians who provided clinical and genomic information. We extend our sincere appreciation to the patient and his family for their participation in this study. We thank Lili Mantcheva for assistance with IRB requirements, participant consent, family communication, and sample collection. We also thank Pattrick Gellispie for assistance with the logistics and coordination related to the KCNA7 plasmid.

## Supplementary material

Supplementary material includes description of 3D modeling of KCNA7 missense variant; Figure S1, Figure S2, Figure S3, Figure S4 and Supplementary Table 1.

## List of Members of the Undiagnosed Diseases Network (Version: 10.24.2024)

Jose Abdenur, Maria T. Acosta, David R. Adams, Ben Afzali, Ali Al-Beshri, Eric Allenspach, Aimee Allworth, Raquel L. Alvarez, Justin Alvey, Ashley Andrews, Beatriz Anguiano, Euan A. Ashley, Sanaz Attaripour, Suha Bachir, Carlos A. Bacino, Guney Bademci, Ashok Balasubramanyam, Dustin Baldridge, Erin E. Baldwin, Elsa Balton, Michael Bamshad, Deborah Barbouth, Rebekah Barrick, Donald Basel, Pinar Bayrak-Toydemir, Taylor Beagle, Alan H. Beggs, Edward Behrens, Megan Bell, Hugo J. Bellen, Paul Berger, Jonathan A. Bernstein, Gerard T. Berry, Stephanie Bivona, Lauren Blieden, Elizabeth Blue, Devon Bonner, Brett Bordini, Nicholas Borja, Lorenzo Botto, Steven Boyden, Lauren C. Briere, Elizabeth A. Burke, Lindsay C. Burrage, Francisco Bustos, Manish J. Butte, Russell Butterfield, Peter Byers, William E. Byrd, Kaitlin Callaway, John Carey, George Carvalho, Thomas Cassini, Chun-Hung Chan, Sirisak Chanprasert, Elizabeth C. Chao, Hsiao-Tuan Chao, Ivan Chinn, Gary D. Clark, Terra R. Coakley, Laurel A. Cobban, Joy D. Cogan, Matthew Coggins, F. Sessions Cole, Erin Conboy, Brian Corner, Rosario I. Corona, William J. Craigen, Andrew B. Crouse, Vishnu Cuddapah, Hongzheng Dai, Nitsuh K. Dargie, Kahlen Darr, Surendra Dasari, Joie Davis, Margaret Delgado, Esteban C. Dell’Angelica, Patricia Dickson, Katrina Dipple, Naghmeh Dorrani, Jessica Douglas, Precilla D’Souza, Filippo Pinto e Vairo, Abdul Elkadri, Sara Emami, Lisa T. Emrick, Christine M. Eng, Cecilia Esteves, Kimberly Ezell, Layal F. Abi Farraj, Elizabeth L. Fieg, Paul G. Fisher, Brent L. Fogel, Jiayu Fu, William A. Gahl, Rebecca Ganetzky, Emily Glanton, Ian Glass, Page C. Goddard, Joanna M. Gonzalez, John E. Gorzynski, Brett H. Graham, Andrea Gropman, Meghan C. Halley, Rizwan Hamid, Neil Hanchard, Kelly Hassey, Frances High, Fuki M. Hisama, Ingrid A. Holm, Jason Hom, Martha HorikePyne, Alden Huang, Yan Huang, Monika Weisz Hubshman, Anna Hurst, Wendy Introne, Gail P. Jarvik, Orpa Jean-Marie, Tanner D. Jensen, Vaidehi Jobanputra, Oguz Kanca, Yigit Karasozen, Laura Keehan, Shamika Ketkar, Dana Kiley, Gonench Kilich, Eric Klee, Shilpa N. Kobren, Isaac S. Kohane, Jennefer N. Kohler, Bruce Korf, Susan Korrick, Elijah Kravets, Runjun Kumar, Albert R. La Spada, Seema R. Lalani, Brendan C. Lanpher, Ian R. Lanza, Kumarie Latchman, Kimberly LeBlanc, Brendan H. Lee, Miranda Leitheiser, Kathleen A. Leppig, Mia Levanto, Richard A. Lewis, Rachel Li, Khurram Liaqat, Pengfei Liu, Nicola Longo, Joseph Loscalzo, Richard L. Maas, Ellen F. Macnamara, Calum A. MacRae, Valerie V. Maduro, Audrey Stephannie C. Maghiro, Rachel Mahoney, May Christine V. Malicdan, Tarun K. K. Mamidi, Lili Mantcheva, Rong Mao, Ronit Marom, Gabor Marth, Beth A. Martin, Martin G. Martin, Julian A. Martínez-Agosto, Shruti Marwaha, Taylor Maurer, Julie McCarrier., Allyn McConkie-Rosell, Ashley McMinn, Erin McRoy, Hector Rodrigo Mendez, Matthew Might, Mohamad Mikati, Alexander Miller, Danny Miller, Ghayda Mirzaa, Breanna Mitchell, Stephen B. Montgomery, Paolo Moretti, Jennifer Morgan, Marie Morimoto, Tahseen Mozaffar, John J. Mulvihill, Lindsay Mulvihill, Michael Muriello, Mariko Nakano-Okuno, Stanley F. Nelson, Serena Neumann, Thomas J. Nicholas, Donna Novacic, Devin Oglesbee, James P. Orengo, Rebecca Overbury, Laura Pace, Stephen Pak, J. Carl Pallais, Neil H. Parker, Alex Paul, LeShon ’ Peart, Leoyklang Petcharet, John A. Phillips III, Jennifer E. Posey, Lorraine Potocki, Aaron Quinlan, Daniel J. Rader, Ramakrishnan Rajagopalan, Deepak A. Rao, Anna Raper, Wendy Raskind, Adriana Rebelo, Chloe M. Reuter, Lynette Rives, Lance H. Rodan, Martin Rodriguez, Jill A. Rosenfeld, Elizabeth Rosenthal, Francis Rossignol, Bianca E. Russell, Marla Sabaii, Mohamad Saifeddine, Jacinda B. Sampson, Suzanne Sandmeyer, Timothy Schedl, Jason Schend, Lisa Schimmenti, Kelly Schoch, Jennifer Schymick, Daryl A. Scott, Elaine Seto, Vandana Shashi, Emily Shelkowitz, Sam Sheppeard, Jimann Shin, Amanda M. Shrewsbury, Edwin K. Silverman, Kathyrn Singh, Giorgio Sirugo, Kathy Sisco, Tammi Skelton, Cara Skraban, Carson A. Smith, Kevin S. Smith, Jared Sninsky, Lilianna SolnicaKrezel, Ben Solomon, Rebecca C. Spillmann, Maija-Rikka Steenari, Andrew Stergachis, Joan M. Stoler, Kathleen Sullivan, Shamil R. Sunyaev, David A. Sweetser, Barbara N. Pusey Swerdzewski, Virginia Sybert, Holly K. Tabor, Queenie Tan, Arjun Tarakad, Herman Taylor, Mustafa Tekin, Willa Thorson, Cynthia J. Tifft, Camilo Toro, Alyssa A. Tran, Kayla M. Treat, Brianna Tucker, Rachel A. Ungar, Adeline Vanderver, Andres Vargas, Matt Velinder, James Verbsky, Francesco Vetrini, Eric Vilain, Dave Viskochil, Tiphanie P. Vogel, Colleen E. Wahl, Melissa Walker, Nicole M. Walley, Jennifer Wambach, Michael F. Wangler, Alistair Ward, Isum Ward, Patricia A. Ward, Stephanie M. Ware, Daniel Wegner, Corrine K. Welt, Mark Wener, Monte Westerfield, Matthew T. Wheeler, Jordan Whitlock, Laurens Wiel, Brandon M. Wilk, Lynne A. Wolfe, Heidi Wood, Kim Worley, Elizabeth A. Worthey, Changrui Xiao, Shinya Yamamoto, Michael Zimmermann, Stephan Zuchner. L.

