## Supplementary material for "A New Disease Gene for Hypokalemic Periodic Paralysis, *KCNA7*, Established in a Multigenerational Family"

### Supplemental Material

### 3D modeling of the missense *KCNA7* variant

Computational analysis revealed that the structural conformation and function of voltage-gated potassium channel (Kv1.7) protein might be substantially altered by the substitution of Serine (S), a polar uncharged residue, at the position of arginine (R), a positively charged amino acid (Fig. S1A-B). We further assessed the polar contacts of the residue 278 using the molecular visualization tool PyMol. In the wild-type protein, ARG 278 forms strong hydrogen bonds with the neighboring Leu274 (3.1Å), Ala275 (3.2 Å), Arg281 (3.3 Å), Leu282 (3.4 Å) (Fig. S1C). In the mutant protein, Ser278 forms strong hydrogen bonds with the neighboring Leu274 (3.1 Å, 3.5 Å), Ala275 (3.3 Å), Leu282 (3.5 Å) (Fig. S1D). The mutation reduces the number and strength of hydrogen bonds, especially the loss of interaction with Arg281 as well as slightly increasing bond distances and weakening stability (Fig. S1E). DynaMut analysis predicts a stability change of (ΔΔGStability) = -0.46 kcal/mol, indicating destabilization due to loss of hydrophobic core packing that disrupts local charge-based interactions, consequently decreasing the local stability. The substitution of Arg278Ser is particularly significant as it occurs in the transmembrane segment 4 helix of the transmembrane domain, a voltage-sensing domain that has frequently repetitive positively charged arginine residues responsible for sensing voltage. Serine substitution might alter secondary structure, affecting protein's helical integrity, disrupting gating-charge transfer and impairing channel openings (Fig. S1F).

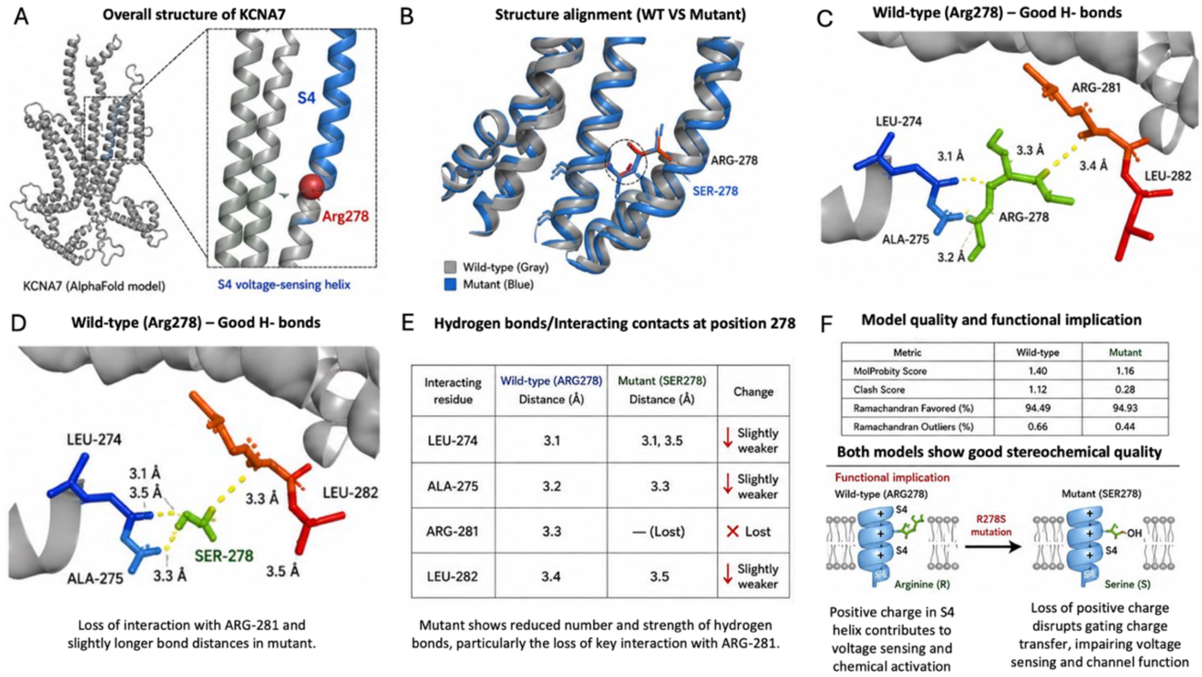


**Figure S1:** Structural and functional impact of the p.Arg278Ser variant in KCNA7. **(A)** Overall structure of KCNA7 based on the AlphaFold model shown in cartoon representation. The boxed region highlights the S4 voltage-sensing helix, and the inset shows the position of Arg278 (red sphere). **(B)** Structural alignment of wild-type (gray) and p.Arg278Ser mutant (blue) proteins. Superposition of the S4 helices indicates that the overall helical conformation is preserved, with localized differences around residue 278. **(C)** Hydrogen bonding interactions in the wild-type protein. Arg278 forms stabilizing hydrogen bonds (yellow dashed lines) with neighboring residues LEU274 (3.1 Å), ALA275 (3.2 Å), ARG281 (3.3 Å), and LEU282 (3.4 Å). **(D)** Hydrogen bonding interactions in the mutant protein. Ser278 maintains interactions with LEU274 (3.1–3.5 Å), ALA275 (3.3 Å), and LEU282 (3.5 Å), but the interaction with ARG281 is lost. **(E)** Quantitative comparison of hydrogen bond interactions at position 278. The mutant shows increased bond distances and loss of interaction with ARG281, indicating a reduction in the number and strength of stabilizing contacts. **(F)** Model quality assessment and functional implications. Both wild-type and mutant models exhibit good stereochemical quality based on MolProbity score, clash score, and Ramachandran statistics. The schematic illustrates that substitution of positively charged arginine with neutral serine in the S4 helix disrupts gating charge transfer, impairing voltage sensing and channel function.

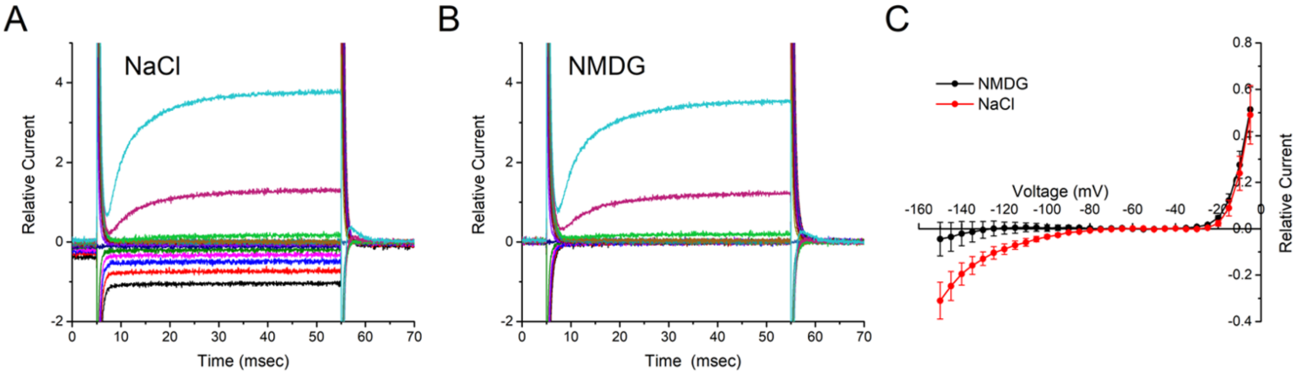


**Figure S2**. The Arg278Ser gating pore in not permeable to NMDG. (A) Representative currents recorded from an oocyte expressing Arg278Ser show inward gating pore currents in a NaCl bath. (B) Currents recorded from the same oocyte, now bathed in NDMG, do not have a detectable inward gating pore component. (C) The mean steady-state current – voltage relationship for oocytes expressing Arg278Ser show inward rectification beginning at -70 mV in NaCl but not NMDG. Data points show the mean ± SEM for the same 4 oocytes bathed in NaCl and then NMDG.

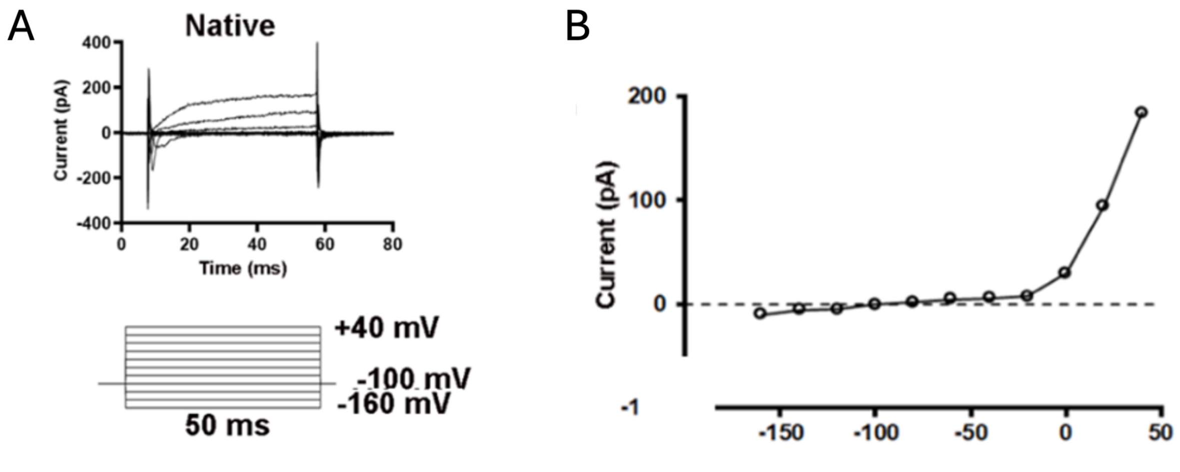


**Figure S3.** Ionic currents in native HEK293 cells. **A**, Exemplar whole-cell current traces for a native HEK293 cell. Currents were elicited by 50-ms voltage commands ranging from -150 mV to +40 mV in 20-mV steps. Holding potential was -100 mV (voltage-clamp protocol is shown in the lower panel). Interval between voltage commands was 2 s. Traces are representative of 3 cells. **B**, Average steady-state current density – voltage relations for native HEK293 cells. Currents were averaged across the terminal 10 ms of each 50-ms pulse and plotted as a function of command voltage. Currents were not corrected for unspecific leak. Values are mean ± SEM.

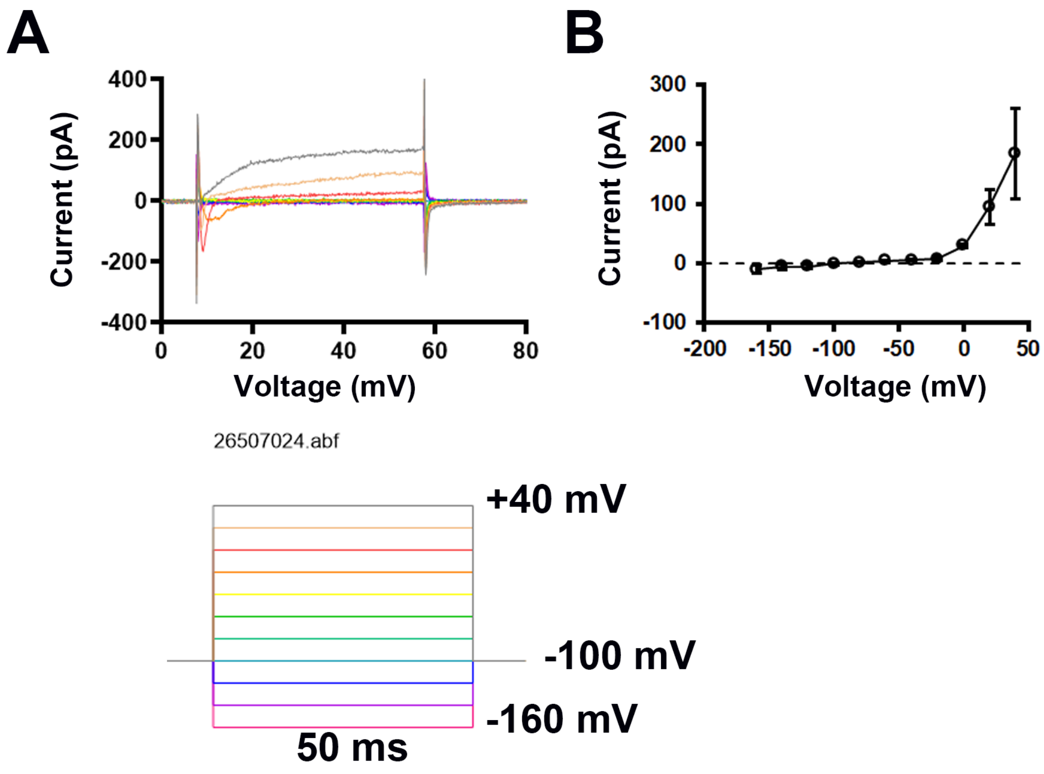


**Figure S4** Tail current protocol to measure the voltage dependence of channel activation. Representative currents are shown for recordings from HEK cells expressing WT or Arg278Ser KV1.7 channels. The relative peak current at the onset of the "tail" potential at -40 mV (end of 500 msec voltage step) was measured to determine the relative channel activation during the preceding test pulse as shown in Figure 5E.

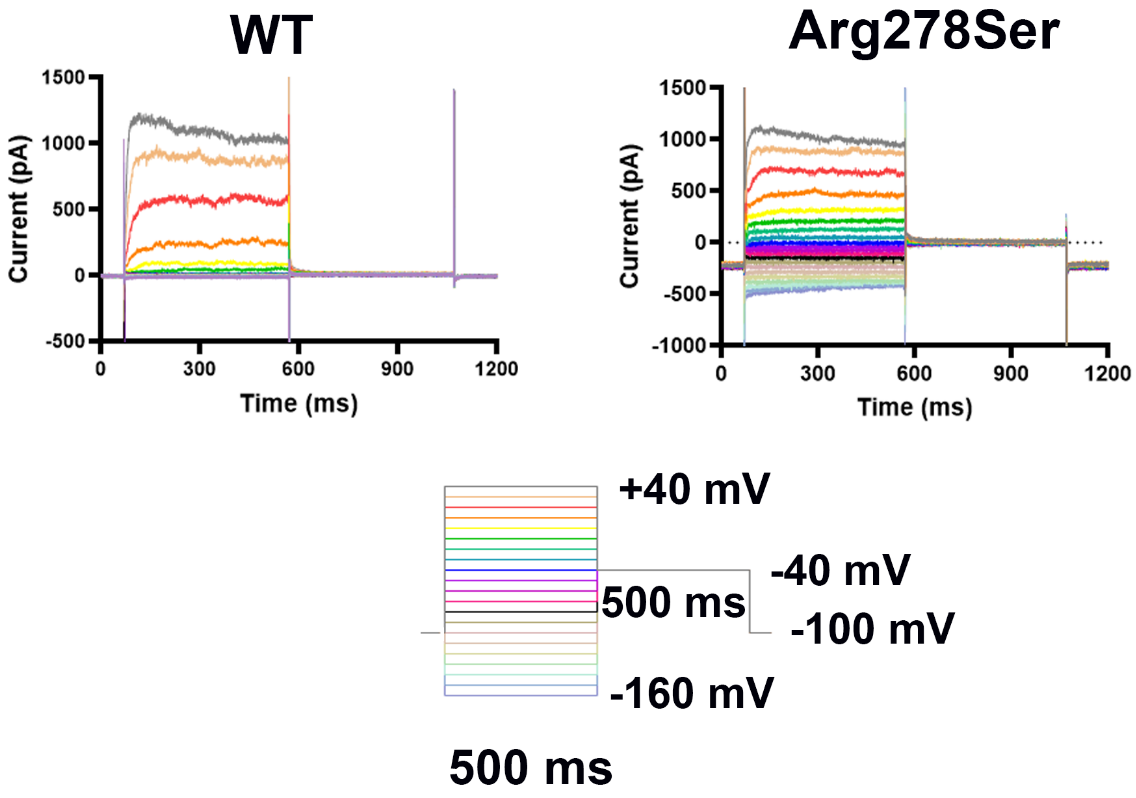

WT
Arg278Ser
Current (pA)
1500
1000
500
0
-500
-1000
0
300
600
900
1200
Time (ms)
+40 mV
-40 mV
500 ms
-100 mV
-160 mV
500 ms

**Supplementary Table 1:** In silico pathogenicity predictions and population frequency of the KCNA7 missense variant (p.Arg278Ser).

| | |
|---|---|
| Gene | *KCNA7* (NM_031886.3) |
| Variant | c.834A>C (p.Arg278Ser) |
| Genomic coordinates (hg19) | chr19:49573857 |
| **In silico Prediction** | |
| SIFT | Deleterious |
| Mutation Taster | Disease causing |
| Polyphen2 | Deleterious |
| CADD | 21.1 |
| REVEL | 0.846 |
| **Population Summary** | |
| gnomAD (v3.1 and v4.1) Allele Frequency | Absent |
| All of US (Allele Frequency) | 0.000004 |
